# Green function of correlated genes and the mechanical evolution of protein

**DOI:** 10.1101/246082

**Authors:** Sandipan Dutta, Jean-Pierre Eckmann, Albert Libchaber, Tsvi Tlusty

## Abstract

There has been growing evidence that cooperative interactions and configurational rearrangements underpin protein functions. But in spite of vast genetic and structural data, the information-dense, heterogeneous nature of protein has held back the progress in understanding the underlying principles. Here we outline a general theory of protein that quantitatively links sequence, dynamics and function: The protein is a strongly-coupled amino acid network whose interactions and large-scale motions are captured by the mechanical propagator, also known as the Green function. The propagator relates the gene to the connectivity of the amino acid network and the transmission of forces through the protein. How well the force pattern conforms to the collective modes of the functional protein is measured by the fitness. Mutations introduce localized perturbations to the propagator which scatter the force field. The emergence of function is manifested by a *topological transition* when a band of such perturbations divides the protein into subdomains. *Epistasis* quantifies how much the combined effect of multiple mutations departs from additivity. We find that epistasis is the nonlinearity of the Green function, which corresponds to a sum over multiple scattering paths passing through the localized perturbations. We apply this mechanical framework to the simulations of protein evolution, and observe long-range epistasis which facilitates collective functional modes. Our model lays the foundation for understanding the protein as an evolved state of matter and may be a prototype for other strongly-correlated living systems.

---

While there are essential differences among the diverse biological functions of proteins, there is also growing recognition that a common physical basis for these functions is the emergence of large-scale, collective patterns of forces and coordinated displacements (1–4). Though the link between protein mechanics and its function is evident and known in certain important mechanisms, such as allostery and induced fit, a basic physical theory that relates sequence evolution to amino acid interactions is still lacking. This is due to the unique nature of this phase of matter: protein is synthesized from twenty distinct species of building blocks, whose arrangement is selected via a long evolutionary process, rendering it much more heterogeneous and non-random than conventional phases of matter (5–9).

The present paper introduces a physical theory that treats protein as an amorphous amino acid network. Mutations that substitute one amino acid by another tweak the interactions, allowing the network to evolve a mechanical function: in response to a localized force, the protein will undergo a large-scale conformational change. This models global structural transitions that are essential to the functions of binding, catalysis and molecular recognition in numerous proteins (10–14). For example, many enzymes are known to deform upon binding to their targets (15–17) by hinge-like rotations, twists or shear-like sliding (18, 19). Recent works link such large-scale motions with the emergence of a marginally stable ‘shear band’ or ‘channel’ in the protein (18–23), and other works highlighted the role of such collective interactions in allostery, the transduction of signal between distant sites in the protein (24–30). Previously, we examined the shear band as a mechanism for allostery, and our simple model distilled certain basic features of the genotype-to-phenotype map (22, 23).

Our aim here is different: to construct a physical theory which treats protein in terms of evolving condensed matter, and thus gives formal mechanical interpretation to the basic determinants of evolution. Moreover, the evolution of a ‘channel’ (18, 23, 31) is a general phenomenon, more basic than allostery. Channels may first evolve as part of a catalytic or molecular recognition mechanism, where the binding of a ligand at one end of a channel may trigger a global motion of an enzyme (Fig. 1A,B). Allostery may emerge later, when the other end of the channel evolves affinity to another ligand.

**Fig. 1.**
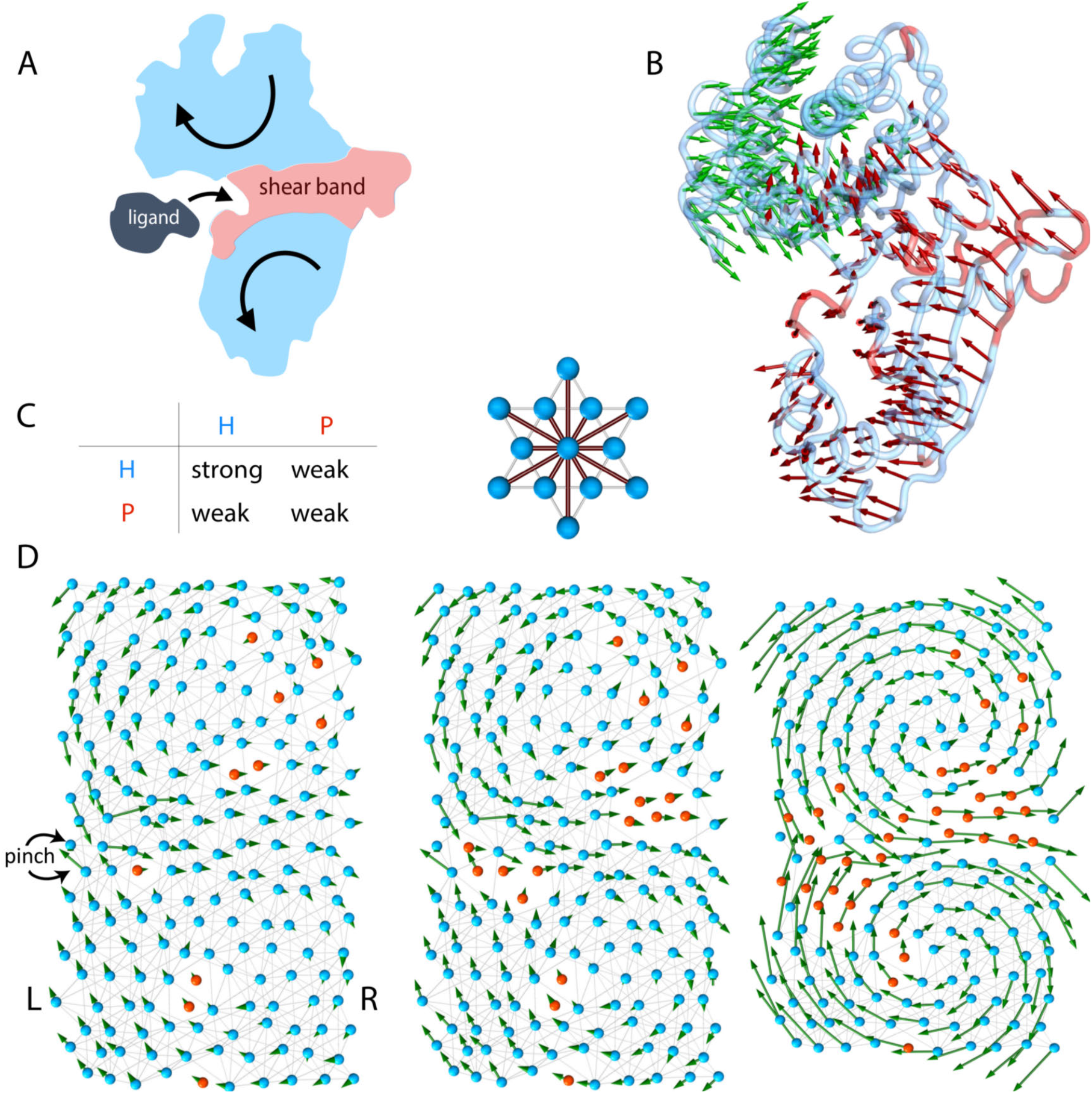
Protein as an evolving machine and propagation of mechanical forces. **(A)** The cartoon depicts the formation of a softer ‘shear band’ (red) separating the protein into two rigid subdomains (light blue). This topology facilitates large-scale conformational transitions. When a ligand binds, the biochemical function involves a low-energy, hinge-like or shear motion (arrows). (B) Shear band and large scale motion in a real protein: the arrows show the displacement of all amino acids (AAs) in human glucokinase when it binds glucose (PDB structures: 1v4s and 1v4t). We calculated the shear field using the method of (18, 32). The coloring shows a high shear region (red) separating two low-shear domains that move as rigid bodies (similar to A). **The mechanical model**. (C) The protein is made of two species of AAs, polar (P, red) and hydrophobic (H, blue) whose sequence is encoded in a gene. Each amino acid forms weak or strong bonds with its 12 near neighbors (right) according to the interaction rule in the table (left). (D) The protein is made of 20 *×* 10 = 200 AAs whose positions are randomized from a regular triangular lattice. Strong bonds are shown as gray lines and weak bonds are not shown for clarity. Evolution begins from a random configuration (left) and evolves by mutating one AA at each step, switching between H and P. The fitness is the mechanical response to a localized force probe (‘pinch’), which is calculated from the Green function (see text). Mutations that amplify the fitness are accepted and the rest are rejected. After about 10^3^ mutations, and about a dozen of accepted beneficial mutations (middle: intermediate stage), the evolution finds a solution (right). The green arrows show the mechanical response: a hinge-like, low-energy motion with a shear band starting at the probe and traversing the protein, qualitatively similar to the displacement field and shear band shown in glucokinase (B). The response to the pinch in the initial, random protein is minute (left) and is amplified by evolving the connectivity of the AA network (middle) until a functional collective mode appears (right).

We treat protein as a strongly-coupled many-body system: an amino acid (AA) network whose topology is determined by the gene, which evolves to exhibit a specific large-scale dynamic mode (Fig. 1C,D). Gene sequences encode chains of AAs of two species(33), hydrophobic (H) and polar (P). The chain is folded and the AAs interact with their nearest neighbors, through strong and weak bonds according their chemical nature. We show that the application of a Green function (34) – which is widely used to study inanimate condensed matter (35) – is a natural way to understand the many-body physics of living matter. The Green function measures how the protein responds to a localized impulse, as triggered by a ligand docking at a binding site, via propagation of forces and motion. It allows us to identify and focus on the relevant excitations in the protein, its collective degrees of freedom. The propagation of mechanical response across the protein defines its fitness and directs the evolutionary search.

### Significance Statement

Much is known about the structure, genetics and function of proteins. But a theory that relates the three is still missing. This is because protein is a unique phase of matter: A folded hetero-polymer made of twenty species of amino acids, whose interactions are both cooperative and weak. Moreover, protein evolves to perform biological tasks, and is therefore non-random and information-rich. We present a physical theory that captures these hallmarks of protein in terms of the Green function, which directly links the gene to force propagation and collective dynamics in the protein. The Green function framework allows us to derive simple mechanical expressions for central biological quantities, which are often hard to formulate analytically, in particular fitness, mutations and epistasis.

Thus, the Green function explicitly defines the map: gene → amino acid network → protein dynamics → function. We use this map to quantify the effect of mutation and to examine *epistasis*, the interaction among mutations, which gives rise to the departure of the fitness function from additivity (36–38). Growing evidence suggests that epistasis is a major determinant of the protein’s fitness landscape (39–42). But due to the complexity of the underlying physical interactions, epistasis remains elusive and hard to deduce. Because a mutation perturbs the Green function, and as a result deflects and scatters the propagation of force through the protein. This allows us to quantify epistasis, even among spatially distant mutations, in terms of “multiple scattering" pathways. These indirect physical interactions appear as long-range correlation in the co-evolution of protein sequences.

The main findings are:

1. Using a Metropolis-type evolution algorithm, solutions are quickly found, typically after ∼10^3^ steps. Mutations add localized perturbations to the AA network, which are eventually arranged by evolution into a continuous shear band. The emergence of protein function is signaled by a topological transition which occurs when this shearable band of weakly connected amino acids separates the protein into two rigid subdomains. This allows the appearance of a low-energy, functional mode thanks to the softness of the band.
2. The set of solutions is sparse: there is a huge reduction of dimension between the space of genes to the spaces of force and displacement fields, which are constrained by the mechanical function.
3. There is a tight correspondence between the correlations in gene sequences and those in the mechanical fields. In particular, epistasis among codons in the gene originates from the mechanical interactions among the corresponding AAs in the protein.

## Model: Protein as an evolving machine

### The amino acid network and its Green function

To examine the large-scale, slow dynamics of protein, we use a coarse grained description in terms of an elastic network (43–45) whose connectivity and interactions are encoded in a gene (Fig. 1C,D). The protein is a chain of *n_a_* = 200 amino acids, *a_i_* (*i* = 1*, …, n_a_*) folded into a 10 × 20 two-dimensional hexagonal lattice (*d* = 2). We follow the simplified HP model (33) with two species of AAs, hydrophobic (*a_i_* = H) and polar (*a_i_* = P). The AA chain is encoded in a gene **c**, a sequence of 200 binary codons, where *c_i_* = 1 encodes a H AA and *c_i_* = 0 encodes a P. We consider a constant fold, so any particular codon *c_i_* in the gene encodes an AA *a_i_* at a certain constant position **r**_*i*_ in the protein. The positions **r**_*i*_ are randomized to make the network amorphous. These *n_d_* = *d · n_a_* = 400 degrees-of-freedom are stored in a vector **r**. Except the ones at the boundaries, every AA is connected by harmonic springs to *z* = 12 nearest and next-nearest neighbors. There are two flavors of bonds according to the chemical interaction which is defined as an AND gate: a strong H−H bond and weak H−P and P−P bonds. The strength of the bonds determines the mechanical response of the network to a displacement field **u**, when the AAs are displaced as **r**_*i*_ → **r**_*i*_ + **u**_*i*_. The response is captured by Hooke’s law that gives the force field **f** induced by a displacement field, **f** = **H**(**c**) **u**. The analogue of the spring constant is the Hamiltonian **H**(**c**), a *n_d_×n_d_* matrix, which records the connectivity of the network and the strength of the bonds. **H**(**c**) is a *nonlinear* function of the gene **c**, reflecting the AA interaction rules of Fig. 1C (see [11], Methods).

The fitness of the protein is related to the *inverse problem* to Hooke’s law: given a localized force **f** (‘pinch’), evolution searches for a protein which will respond by a prescribed large-scale motion **u**. In induced fit, for example, specific binding of a substrate should induce global deformation of an enzyme. The response is determined by the *Green function* **G** (34), 
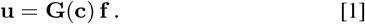

**G** is the mechanical propagator that measures the transmission of signals from the force source **f** across the protein (Fig. 2A). Equation [1] constitutes an explicit *genotype-to-phenotype map* from the genotype **c** to the mechanical phenotype **u**: **c** → **u**(**c**) = **G**(**c**)**f**. This reflects the dual nature of the Green function **G**: in the phenotype space, it is the mechanical propagator which turns a force into motion, **u** = **Gf**, whereas it is also the nonlinear function which maps the gene into a propagator, **c** → **G**(**c**).

**Fig. 2.**
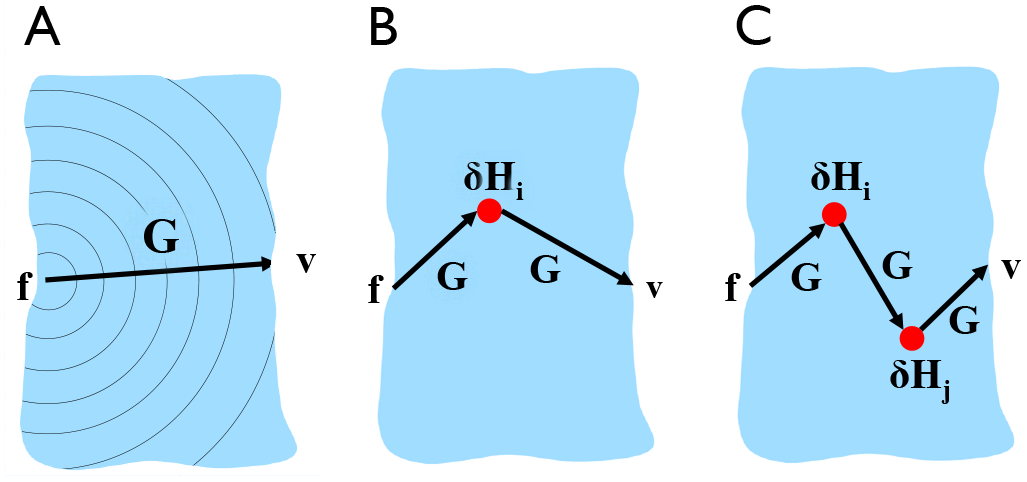
Force propagation, mutations and epistasis. **(A)** The Green function **G** measures the propagation the mechanical signal across the protein (blue) from the force source **f** (pinch) to the response site **v**, depicted as a “diffraction wave". (B) A mutation *δ***H**_*i*_ deflects the propagation of force. The effect of the mutation on the propagator *δ***G** can be described as a series of multiple scattering paths [6]. The diagram shows the first scattering path, **G** *δ***H**_*i*_**G**. (C) Epistasis is the departure from additivity of the combined fitness change of two mutations. The epistasis between two mutations, *δ***H**_*i*_ and *δ***H**_*j*_, is equivalent to a series of multiple scattering paths[9]. The diagram shows the path **G** *δ***H**_*i*_ **G** *δ***H**_*j*_ **G**.

When the protein is moved as a rigid body, the lengths of the bonds do not change and the elastic energy cost vanishes. A 2D protein has *n*_0_ = 3 such zero modes (Galilean symmetries), two translations and one rotation, and **H** is therefore always singular. Hence, Hooke’s law and [1] imply that **G** is the pseudo-inverse, **G**(**c**) = **H**(**c**)^+^ (46, 47), which amounts to inversion of **H** in the non-singular sub-space of the *n_d_ − n*_0_ = 397 non-zero modes (Methods). A related quantity is the resolvent, **G**(*ω*) = (*ω* − **H**)^−1^, whose poles are at the energy levels of **H**, *ω* = λ*_k_*.

The fitness function drives the evolution of the shear band by rewarding strong mechanical response to a localized probe (‘pinch’, Fig. 1D). This probe is a force dipole at two neighboring AAs, *p*′ and *q*′ on the left side of the protein (L), **f**_*q*′_ = −**f**_*p*′_. The desired motion is specified by a displacement vector **v**, with a ‘dipole’ response, **v**_*q*_ = −**v**_*p*_, on the right side of the protein (R). The protein is fitter if the pinch **f** produces a large deformation in the direction specified by **v**, which indicates that the protein is divided into two rigid subdomains moving along a softer shear band. To this end, we evolve the AA network to increase a *fitness function F*, which is the projection of the displacement **u** = **Gf** on the prescribed response **v**, 
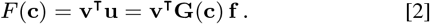

This highlights the role of the Green function **G** in driving the evolution process: Equation [2] defines the *fitness landscape F* (**c**), which is the inner product of **G** between the pinch and the desired response. Here we examine particular examples for a localized ‘pinch’ **f** and desired response **v**, which drive the emergence of a hinge-like mode.However, the present Green function approach is general and can as well treat more complex, multi-site force patterns and multi-domain dynamical modes.

### Evolution searches in the mechanical fitness landscape

Our simulations search for a shear band that starts at a specific site on the L side (pinch) but may end at any of the sites on the R side. This gives rise to a wider shear band that allows the protein to perform more general mechanical tasks (in contrast to the specific allostery problem). In view of this, we define the fitness as the maximum of *F* [2] over all potential locations of the channel’s output (typically 8−10 sites, Methods). We also tested the model when the band connects the pinch site to a specific site on the other side.

The protein is evolved via a point mutation process where, at each step, we flip a randomly selected codon between 0 and 1. This corresponds to exchanging H and P at a random position in the protein, thereby changing the bond pattern and the elastic response by softening or stiffening the AA network. Evolution starts from a random protein configuration, encoded in a random gene. Typically, we take a small fraction of AA of type P (about 5%), randomly positioned within a majority of H. (Fig. 1D). The high fraction of strong bonds renders the protein stiff, and therefore of low initial fitness *F ≃* 0. At each step, we calculate the change in the Green function *δ***G** (by a method explained below) and use it to evaluate from [2] the fitness change *δF*, 
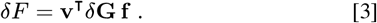

The fitness change *δF* determines the fate of the mutation: we accept the mutation if *δF* ≥ 0, otherwise the mutation is rejected. Since fitness is measured by the criterion of strong mechanical response, it induces softening of the AA network.

The typical evolution trajectory takes about 10^3^ steps. Most are neutral mutations (*δF* ≃ 0) and deleterious ones (*δF <* 0); the latter are rejected. About a dozen or so beneficial mutations (*δ F* > 0) drive the protein towards the solution (Fig. 3A). The increase in the fitness reflects the gradual formation of the channel, while the jump in the shear signals the emergence of the soft mode. The first few beneficial mutations tend to form weakly-bonded, P-enriched regions Dutta *et al*. near the pinch site on the L side, and close to the R boundary of the protein. The following ones join these regions into a floppy channel (a shear band) which traverses the protein from L to R. We stop the simulation when the fitness reaches a large positive value *F*_m_ ∼ 5. The corresponding gene **c**_*_ encodes the functional protein. The adhoc value *F*_m_ ∼ 5 signals slowing down of the fitness curve towards saturation at *F* > *F*_m_, as the channel forms and now only continues to widen. In this regime, even a tiny pinch will easily excite a large-scale motion with a distinct high-shear band (Fig. 1D right).

**Fig. 3.**
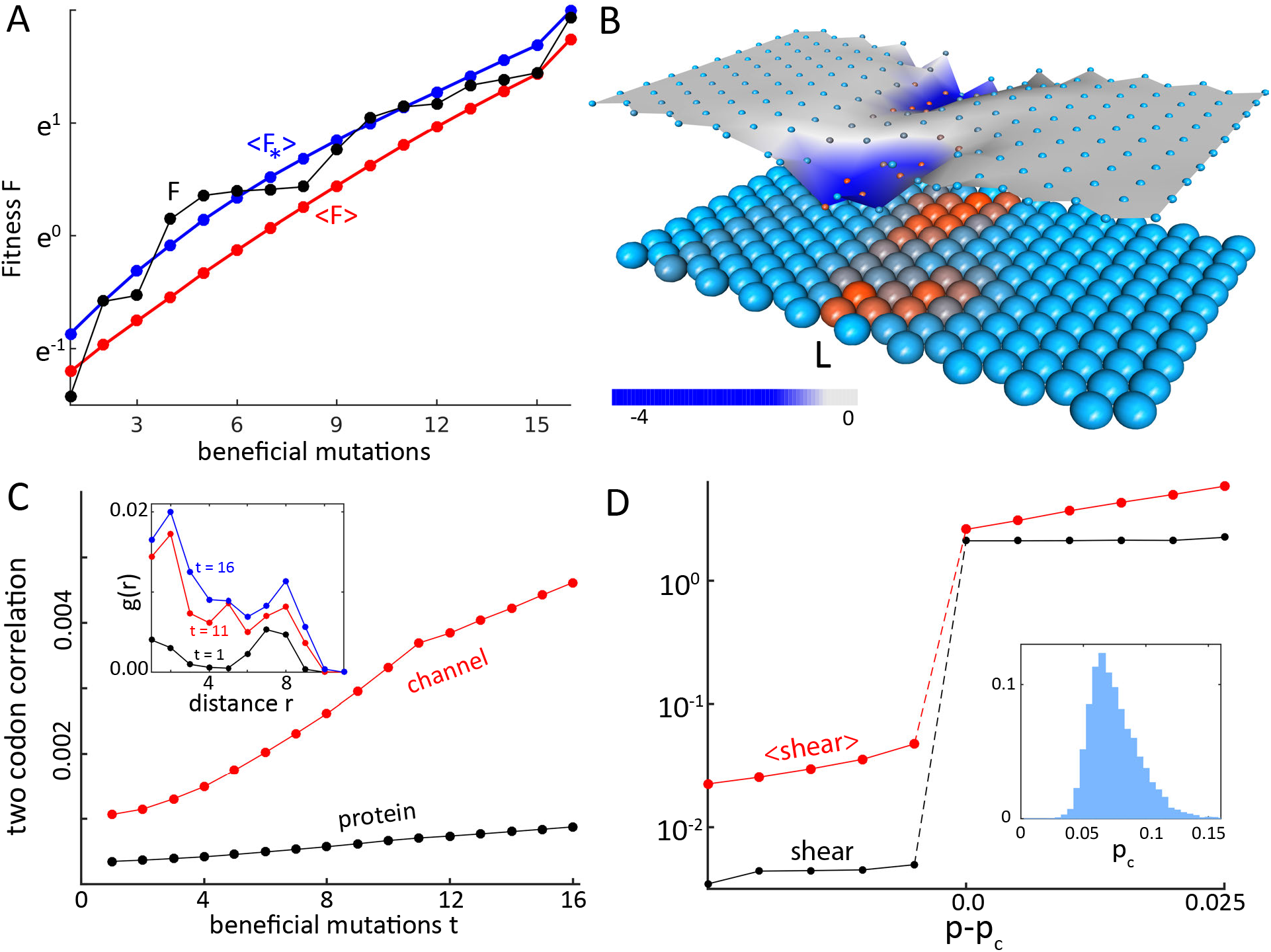
The mechanical Green function and the emergence of protein function. **(A)** Progression of the fitness *F* during the evolution run shown in Fig. 1D (black), together with the fitness trajectory averaged over ∼10^6^ runs 〈*F*〉 (red). Shown are the last 16 beneficial mutations towards the formation of the channel. The contribution of the emergent low-energy mode 〈*F_*_*〉 (blue) dominates the fitness [4]. (B) Landscape of the fitness change *F* [3], averaged over ∼10^6^ solutions, for all 200 possible positions of point mutations at a solution. Underneath, the average AA configuration of the protein is shown in shades of red (P) and blue (H). In most sites, mutations are neutral, while mutations in the channel are deleterious on average. (C) The average magnitude of the two-codon correlation *|Q_ij_|* [5] in the shear band (AAs in rows 7−13, red) and in the whole protein (black) as a function of the number of beneficial mutations, *t*. Inset: profile of the spatial correlation *g*(*r*) within the shear band (*i.e., Q_ij_* as a function of the distance *r* = |**r**_*i*_ − **r**_*j*_|), after *t* = 1, 11, 16 beneficial mutations. Distance between nearest AAs is *r* = 1. (D) The mean shear in the protein in a single run (black) and averaged over ∼10^6^ solutions (red), as a function of the fraction of P amino acids, *p*. The values of *p* are shifted by the position of the jump, *p_c_*. Inset: distribution of *p_c_*.

## Results

### Mechanical function emerges at a topological transition

The hallmark of evolution reaching a solution gene **c**_*_ is the emergence of a new zero energy-mode, **u**_*_, in addition to the three Galilean symmetry modes. Near the solution, the energy of this mode λ_*_ closes the spectral gap, λ_*_ → 0 (Fig. 3A), and **G**(*ω*) has a pole at *ω* = 0. As a result, the emergent mode dominates the Green function, 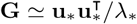. The response to a pinch will be mostly through this *soft mode*, and the fitness [2] will diverge as 
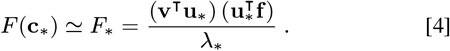

On average, we find that the fitness increases exponentially with the number of beneficial mutations (Fig. 3A). However, beneficial mutations are rare, and are separated by long stretches of neutral mutations. This is evident from the fitness landscape (Fig. 3B), which shows that in most sites the effect of mutations is practically neutral.

The vanishing of the spectral gap, λ_*_ → 0, manifests as a topological transition in the system: the AA network is now divided into two domains that can move independently of each other at low energetic cost. The relative motion of the domains defines the emergent soft mode and the collective degrees-of-freedom, for example the rotation of a hinge or the shear angle.

As the shear band is taking shape, the correlation among codons builds up. To see this, we align the ∼10^6^ simulations and, at each time step, calculate the two-codon correlation *Q_ij_* between all pairs of codons *c_i_* and *c_j_*, 
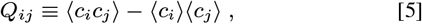
where brackets denote ensemble averages. We find that most of the correlation is concentrated in the forming channel (Fig. 3C), where it is tenfold larger than in the whole protein. In the channel, there is significant long-range correlation shown in the spatial profile of the correlation *g*(*r*) (Fig. 3C, inset).

The soft mode appears at a *dynamical phase transition*, where the average shear in the protein jumps abruptly as the channel is formed and the protein can easily deform in response to the force probe (Fig. 3D). The trajectories are plotted as a function of *p*, the fraction of AAs of type P. The distribution of the critical values *p_c_* is rather wide owing to the random initial conditions and finite-size effects.

### Point mutations are localized mechanical perturbations

A mutation may vary the strength of no more than *z* = 12 bonds around the mutated AA (Fig. 2B). The corresponding perturbation of the Hamiltonian *δ***H** is therefore *localized*, akin to a defect in a crystal (48, 49). The mechanical meaning of mutations can be further explored by examining the perturbed Green function, **G**′ = **G** + *δ***G**, which obeys the Dyson equation (35, 50) (Methods), 
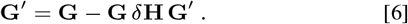

The latter can be iterated into an infinite series, 
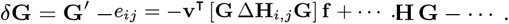

This series has a straightforward physical interpretation as a sum over *multiple scatterings*: As a result of the mutation, the elastic force field is no longer balanced by the imposed force **f**, leaving a residual force field *δ***f** = *δ***H u** = *δ***H G f**. The first scattering term in the series balances *δ***f** by the deformation *δ***u** = **G***δ***f** = **G** *δ***H Gf**. Similarly, the second scattering term accounts for further deformation induced by *δ***u**, and so forth. In practice, we calculate the mutated Green function using the Woodbury formula [12], which exploits the localized nature of the perturbation to accelerate the computation by a factor of ∼10^4^(Methods).

### Epistasis links protein mechanics to genetic correlations

Our mechanical model provides an explicit, calculable definition of epistasis, the deviation from additivity of the combined fitness change of interacting mutations (Fig 2C). We take a functional protein obtained from the evolution algorithm and mutate an AA at a site *i*. This mutation induces a change in the Green function *δ***G**_*i*_ (calculated by [12]) and hence in the fitness function *δF_i_* [3]. One can similarly perform another independent mutation at a site *j*, producing a second deviation, *δ***G**_*j*_ and *δF_j_* respectively. Finally, starting again from the original solution one mutates both *i* and *j* simultaneously, with a combined effect *δ***G**_*i,j*_ and *δF_i,j_*. The epistasis *e_ij_* measures the departure of the double mutation from additivity of two single mutations, 
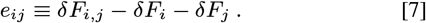

To evaluate the average epistatic interaction among AAs, we perform the double mutation calculation for all 10^6^ solutions and take the ensemble average *E_ij_* = 〈*e_ij_*〉. Landscapes of *E_ij_* show significant epistasis in the channel (Fig. 4). AAs outside the high shear region show only small epistasis, since mutations in the rigid domains hardly change the elastic response. The epistasis landscapes (4A-C) are mostly positive since the mutations in the channel interact antagonistically (51): after a strongly deleterious mutation, a second mutation has a smaller effect.

Definition [7] reveals a direct link between epistasis and protein mechanics: The *nonlinearity* (‘curvature’) of the Green function measures the deviation of the mechanical response from additivity of the combined effect of isolated mutations at *i* and *j*, 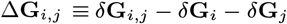. The epistasis *e_ij_* is simply the inner product value of this nonlinearity with the pinch and the response, 
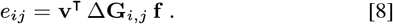

Relation [8] reveals an explicit origin for epistasis from mechanical forces among mutated AAs.

In the gene, epistatic interactions are manifested in codon correlations as shown in Fig. 4D, which depicts two-codon correlations *Q_ij_* from the alignment of ∼10^6^ functional genes **c**_*_ [5]. We find a tight correspondence between the mean epistasis *E_ij_* = 〈*e_ij_*〉 and the codon correlations *Q_ij_*. Both patterns exhibit strong correlations in the channel region with a period equal to channel’s length, 10 AAs. The similarity in the patterns of *Q_ij_* and *E_ij_* indicates that a major contribution to the long-range, strong correlations observed among aligned protein sequences stems from the mechanical interactions propagating through the AA network.

**Fig. 4.**
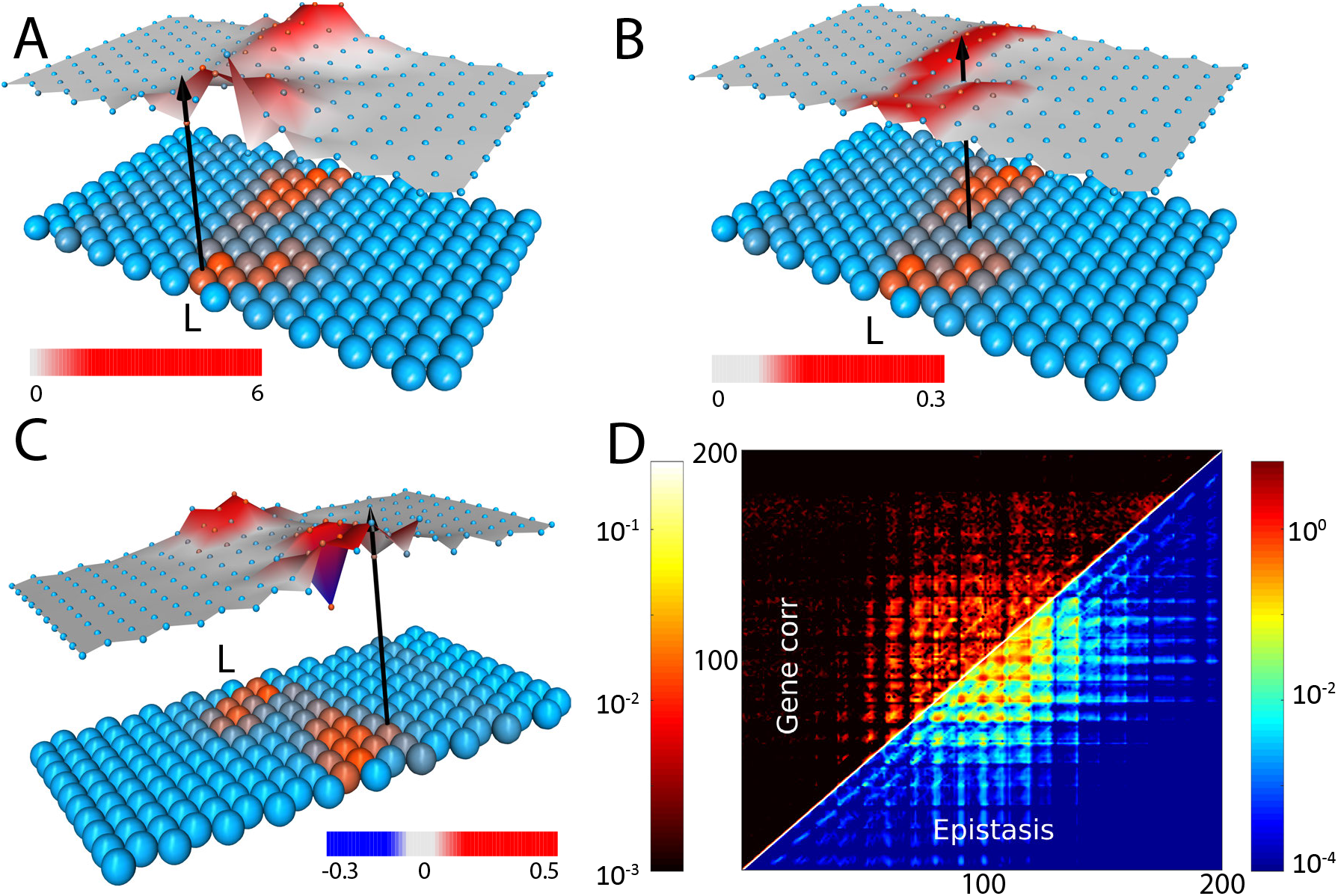
Mechanical Epistasis. The epistasis [7], averaged over ∼ 10^6^ solutions *E_ij_* = 〈*e_ij_*〉, between a fixed AA at position *i* (black arrow) and all other positions *j*. Here, *i* is located at (A) the binding site, (B) the center of the channel, and (C) slightly off the channel. Underneath, the average AA configuration of the protein is drawn in shades of red (P) and blue (H). Significant epistasis mostly occurs along the P-rich channel, where mechanical interactions are long ranged. Though epistasis is predominantly positive, negative values also occur, mostly at the boundary of the channel (C). Landscapes are plotted for specific output site at R. (D) The two-codon correlation function *Q_ij_* [5] measures the coupling between mutations at positions *i* and *j* [5]. The epistasis *E_ij_* and the gene correlation *Q_ij_* show similar patterns. Axes are the positions of *i* and *j* loci. Significant correlations and epistasis occur mostly in and around the channel region (positions ∼70−130, rows 7−13).

### Epistasis as a sum over scattering paths

One can classify epistasis according to the interaction range. Neighboring AAs exhibit *contact epistasis* (52, 53), because two adjacent perturbations, **H**_*i*_ and *δ***H**_*j*_, AND gate of the interaction table (Fig. 1C), 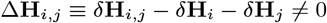 (where *δ***H**_*i,j*_ is the perturbation by both mutations). The leading term in the Dyson series [6] of Δ**G**_*i,j*_ is a single scattering from an effective perturbation with an energy Δ**H**_*i,j*_, which yields the epistasis 
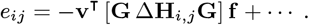

*Long-range epistasis* among non-adjacent, non-interacting perturbations (Δ**H**_*i,j*_ = 0) is observed along the channel (Fig. 4). In this case, [6] expresses the nonlinearity Δ**G**_*i,j*_ as a sum over *multiple scattering* paths which include both *i* and *j* (Fig. 2C), 
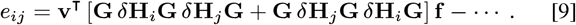

The perturbation expansion directly links long-range epistasis to shear deformation: Near the transition, the Green function is dominated by the soft mode, 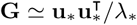, with fitness *F* given by [4]. From [6] and [8] we find a simple expression for the mechanical epistasis as a function of the shear, 
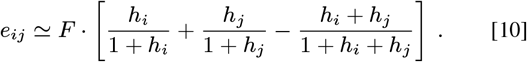

The factor 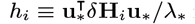 in [10] is the ratio of the change in the shear energy due to mutation at *i* (the expectation value of *δ***H**_*i*_) and the energy λ_*_ of the soft mode, and similarly for *h_j_*. Thus, *h_i_* and *h_j_* are significant only in and around the shear band, where the bonds varied by the perturbations are deformed by the soft mode. When both sites are outside the channel, *h_i_, h_j_* ≪ 1, the epistasis [10] is small, *e_ij_* ≃ 2*h_i_h_j_F*. It remains negligible even if one of the mutations, *i*, is in the channel, *h_j_* ≪ 1 ≪ *h_i_*, and *e_ij_* ≃ *h_j_F*. Epistasis can only be long-ranged along the channel when both mutations are significant, *h_i_, h_j_* ≫ 1, and 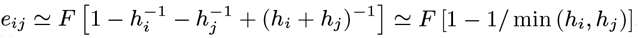. We conclude that epistasis is maximal when both sites are at the start or end of the channel, as illustrated in Fig. 4. The nonlinearity of the fitness function gives rise to antagonistic epistasis since the combined effect of two deleterious mutations is not additive as either mutation is enough to diminish the fitness.

### Geometry of fitness landscape and gene-to-function map

With our mechanical evolution algorithm we can swiftly explore the fitness landscape to reveal its geometry. The genotype space is a 200-dimensional hypercube, whose vertices are all possible genes **c**. The phenotypes reside in a 400-dimensional space of all possible mechanical responses **u**. And the Green function provides the genotypeto-phenotype map [1]. A functional protein is encoded by a gene **c**_*_ whose fitness exceeds a large threshold, 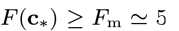, and the functional phenotype is dominated by the emergent zero-energy mode, 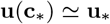 (Fig. 3A). We also characterize the phenotype by the pattern of the shear field **s**_*_ (Methods).

The singular value decomposition (SVD) of the 10^6^ solutions returns a set of eigenvectors, whose ordered eigenvalues show their significance in capturing the data (Methods). The SVD spectra reveal strong correspondence between the genotype **c**_*_ and the phenotype, **u** and **s**_*_ (Fig. 5). In all three data sets, the largest eigenvalues are discrete and stand out from the bulk continuous spectrum. These are the *collective degrees-of-freedom*, which show loci in the gene and positions in the “flow” (*i.e*., displacement) and shear fields that tend to vary together.

**Fig. 5.**
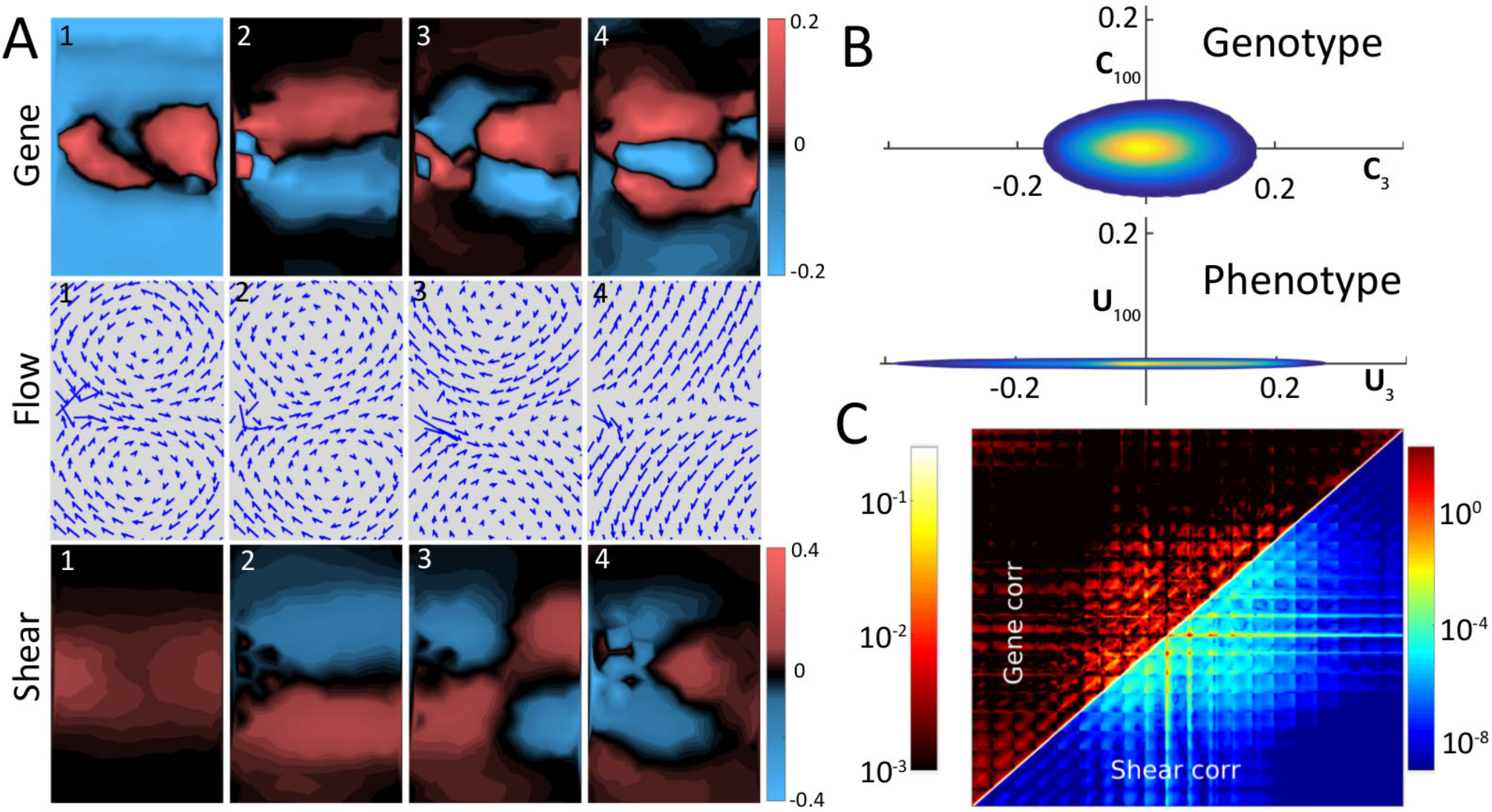
From gene to mechanical function: spectra and dimensions. **(A)** The first four SVD eigenvectors (see text), of the gene **C**_*k*_ (top row), the displacement flow field **U**_*k*_ (middle) and the shear **S**_*k*_ (bottom). (B) Cross sections through the set of solutions in the genotype space (top) and the phenotype space (bottom). Density of solutions is color coded. The genotype cross section is the plane defined by the eigenvectors **C**_3_−**C**_100_, and in the phenotype space by the eigenvectors **U**_3_−**U**_100_ (see text). The dimensional reduction is manifested by the discoid geometry of the phenotype cloud as compared to the spheroid shape of the genotype cloud. (C) Genetic correlations, *Q_ij_* show similarity to correlations in the shear field, **s**_*_ (color coded in log scale).

We examine the correspondence among three sets of eigenvectors, {**U**_*k*_} of the flow, {**C**_*k*_} of the gene, and {**S**_*k*_} of the shear. The first eigenvector of the flow, **U**_1_, is the hinge motion caused by the pinch, with two eddies rotating in opposite directions (Fig. 5A). The next two modes, shear, **U**_2_, and breathing, **U**_3_, also occur in real proteins such as glucokinase (Fig. 1B). The first eigenvectors of the shear **S**_1_ and of the gene sequence **C**_1_ show that the high shear region is mirrored as a P-rich region, where a mechanical signal may cause local rearrangement of the AAs by deforming weak bonds.In the rest of the protein, the H-rich regions move as rigid bodies with insignificant shear. The higher gene eigenvectors, **C**_*k*_ (*k >*1), capture patterns of correlated genetic variations. The striking similarity between the sequence correlation patterns **C**_*k*_ and the shear eigenvectors **S**_*k*_ reveals a tight genotype-to-phenotype map, as is further demonstrated in the likeness of the correlation matrices of the AA and shear flow (Fig. 5C).

In the phenotype space, we represent the displacement field **u** in the SVD basis, {**U**_*k*_} (Fig. 5B). Since ∼90% of the data is explained by the first ∼15 **U**_*k*_, we can compress the displacement field without much loss into the first 15 coordinates. This implies that the set of solutions is a 15-dimensional *discoid* which is flat in most directions. In contrast, representation of the genes **c**_*_ in the SVD frame-of-reference (with the {**C**_*k*_} basis) reveals that in genotype space the solution set is an incompressible 200-dimensional *spheroid* (Fig. 5B). The dramatic dimensional reduction in mapping genotypes to phenotypes stems from the different constraints that shape them (54–57). In the phenotype space, most of the protein is rigid, and only a small number of shear motions are low-energy modes, which can be described by a few degrees-of-freedom. In the genotype space, in contrast, there are many neutral mutations which do not affect the motion of the protein.

## Discussion

Together with DNA and RNA, proteins are the smallest physical objects where the difference between living and inanimate becomes apparent. Understanding proteins is therefore a basic problem of biology, and much effort has been invested in measuring them in great detail. But proteins are highly heterogeneous and dense in information acquired via the evolutionary selection process. A theory of proteins should therefore unify the many-body physics of the amino acid matter with the evolution of genetic information, which together give rise to protein function.

We introduced such a physical theory of protein-like matter, where the mapping between genotype and phenotype takes an explicit form in terms of mechanical propagators (Green functions). As a functional phenotype we take cooperative motion and force transmission through the protein [2]. This allows us to map genetic mutations to mechanical perturbations which scatter the force field and deflect its propagation [3,6] (Fig. 2). The evolutionary process amounts to solving the inverse scattering problem: Given prescribed functional modes, one looks for network configurations, via the mutation process, which yield this low end of the dynamical spectrum.

Epistasis, the interaction among loci in the gene, corresponds to a sum over all multiple scattering trajectories, or equivalently the nonlinearity of the Green function [7,8]. We find that long-range epistasis signals the emergence of a collective functional mode in the protein. The Green function provides a general framework to physically deduce interactions between spatially distant mutations in the protein.

We examined whether a protein’s backbone might affect the results of our model and found that the backbone does not interfere with the emergence of a soft mode whose displacement field is similar to the backbone-less case (Fig. 6). The backbone adds certain features to the gene and deformation field, and might constrain the evolutionary process, but with no impact on the convergence of the search and on the long-range correlations among solutions (Methods).

Overall, the Green function appears as a natural representation of the many-body interactions in the protein. This posits the protein problem in a realm where one can apply the powerful toolbox of condensed matter physics. Some of our simplifying assumptions are inherent to the model, while others can be removed by straightforward generalizations. For example, it is easy to extend the model to 3D geometries, including the configuration of real proteins. Also, one can readily use a more realistic interaction model with twenty AA species (58).

Our model cannot compete with the accuracy and detail of molecular dynamics in simulating real proteins. The aim of our approach is different: to provide an accessible theoretical framework for thinking about basic questions of protein. In particular, the model captures the basic hallmark of protein evolution, the emergence of function as a cooperative pattern of forces and motions. Additionally it highlights how mechanical interactions shape and constrain the progress of evolution.

The Green function proves to a be particularly useful tool for this purpose, and lends itself for intuitive interpretation as a path integral over scattering by mutations. We therefore expect that the present calculable and efficient method will be used in protein evolution models. It may also be applied to study other basic problems in strongly-coupled living matter, such as the evolution of protein interaction networks and molecular recognition.

## Materials and Methods

### The mechanical model of protein

We model the protein as an elastic network made of harmonic springs (43, 45, 59, 60). The connectivity of the network is described by a hexagonal lattice whose vertices are AAs and whose edges correspond to bonds. There are *n_a_* = 10 × 20 = 200 AAs, indexed by Roman letters and *n_b_* bonds, indexed by Greek letters. We use the HP model (33) with two AA species, hydrophobic (*a_i_* = H) and polar (*a_i_* = P). The AA chain is encoded in a gene **c**, where *c_i_* = 1 encodes H and *c_i_* = 0 encodes P, *i* = 1, …, *n_a_*. The degree *z_i_* of each AA is the number of AAs to which it is connected by bonds. In our model, most AA have the maximal degree which is *z* = 12, while AAs at the boundary have fewer neighbors, *z* < 12, see Fig. 1C. The connectivity of the graph is recorded by the adjacency matrix **A**, where **A**_*ij*_ = 1 if there is an edge from *j* to *i*, and **A**_*ij*_ = 0 otherwise. The gradient operator ∇ relates the spaces of edges and vertices (and is therefore of size *n_b_*×*n_a_*): if vertices *i* and *j* are connected by an edge *α*, then ∇_*αi*_ = +1 and ∇_*αj*_ = −1. As in the continuum case, the Laplace operator Δ is the product = Δ = ∇t∇. The non-diagonal elements Δ_*ij*_ are −1 if *i* and *j* are connected and 0 otherwise. The diagonal part Δ of is the degree Δ_*ii*_ = *z_i_*. Hence, we can write the Laplacian as Δ = **Z** − **A**, where **Z** is the diagonal matrix of the degrees *z_i_*.

We embed the graph in Euclidean space 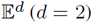, by assigning positions 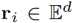 to each AA. We concatenate all positions in a vector **r** of length *n_a_* · d ≡ *n_d_*. Finally, to each bond we assign a spring with constant *k_α_*, which we keep in a diagonal *n_b_×n_b_* matrix **K**. The strength of the spring is determined by the AND rule of the HP model’s interaction table (Fig. 1C), 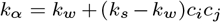 where *c_i_* and *c_j_* are the are the codons of the AA connected by bond α. This implies that a strong H−H bond has *k_α_* = *k_s_*, whereas the weak bonds H−P and P−H have *k_α_* = *k_w_*. We usually take *k_s_* = 1 and *k_w_* = 0.01. This determines a spring network. We also assume that the initial configuration is such that all springs are at their equilibrium length, disregarding the possibility of ‘internal stresses’ (60), so that the initial elastic energy is *ε*_0_ = 0.

We define the ‘embedded’ gradient operator **D** (of size *n_b_×n_d_*) which is obtained by taking the graph gradient ∇, and multiplying each non-zero element (±1) by the corresponding direction vector 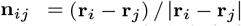 Thus, **D** is a tensor (**D** = Δ_*αi*_**n**_*ij*_), which we store as a matrix (*α* is the bond connecting vertices *i* and *j*). In each row vector of **D**, which we denote as **m**_*α*_ ≡ **D**_*α*,:_, there are only 2*d* non-zero elements. To calculate the elastic response of the network, we deform it by applying a force field **f**, which leads to the displacement of each vertex by **u**_*i*_ to a new position **r**_*i*_ + **u**_*i*_ (see *e.g*., (60)). For small displacements, the linear response of the network is given by Hooke’s law, **f** = **Hu**. The elastic energy is *ε* = **u**^t^**Hu**/2, and the Hamiltonian, **H** = **D**^t^**KD**, is the Hessian of the elastic energy *ε*, 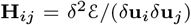 By rescaling, **D** → **K**^½^**D**, which amounts to scaling all distances by 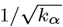, we obtain **H** = **D**^t^**D**. It follows that the Hamiltonian is a function of the gene **H**(**c**), which has the structure of the Laplacian Δ multiplied by the tensor product of the direction vectors. Each *d×d* block **H**_*ij*_ (*i* ≠*j*) is a function of the codons *c_i_* and *c_j_*, 
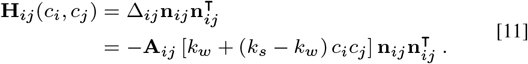

The diagonal blocks complete the row and column sums to zero, 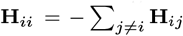

### The inverse problem: Green’s function and its spectrum

The Green function **G** is defined by the inverse relation to Hooke’s law, **u** = **Gf** [1]. If **H** were invertible (non-singular),**G** would have been just **G** = **H**−1. However, **H** is always singular owing to the zero-energy (Galilean) modes of translation and rotation. Therefore, one needs to define **G** as the Moore-Penrose pseudo-inverse (46, 47), **G** = **H**^+^, on the complement of the space of Galilean transformations. The pseudo-inverse can be understood in terms of the spectrum of **H**. There are at least *n*_0_ = *d*(*d* + 1)/2 zero modes: *d* translation modes and *d*(*d* − 1)/2 rotation modes. These modes are irrelevant and will be projected out of the calculation (Note that these modes do not come from missing connectivity of the graph Δ itself but from its embedding in 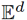). **H** is singular but is still diagonalizable (since it has a basis of dimension *n_d_*), and can be written as the spectral decomposition, 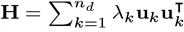, where {λ_*k*_} is the set of eigenvalues and {**u**_*k*_} are the corresponding eigenvectors (note that *k* denotes the index of the eigenvalue, while *i* and *j* denote AA positions). For a non-singular matrix one may calculate the inverse simply as 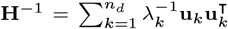. Since **H** is singular we leave out the zero modes and get the pseudo-inverse **H**^+^, 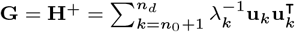. It is easy to verify that if **u** is orthogonal to the zero modes then **u** = **GHu**. The pseudo-inverse obeys the four requirements (46): (i) **HGH** = **H**, (ii) **GHG** = **G**, (iii) (**HG**)^t^ = **HG**, (iv) (**GH**)^t^ = **GH**. In practice, as the projection commutes with the mutations, the pseudo-inverse has most virtues of a proper inverse. The reader might prefer to link **G** and **H** through the heat kernel, 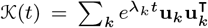. Then, 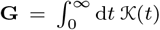 and 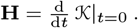.

### Pinching the Network

A pinch is given as a localized force applied at the boundary of the ‘protein’. We usually apply the force on a pair of neighboring boundary vertices, *p*′ and *q*′. It appears reasonable to apply a force dipole,*i.e*., two opposing forces, **f**_*q*′_ = −**f**_*p*′_, since a net force will move the center of mass. This ‘pinch’ is therefore specified by the force vector **f** (of size *n_d_*) whose only 2*d* non-zero entries are **f**_*q*′_ = −**f**_*p*′_. Hence, it has the same structure as a bond vector **m**_*α*_ of a ‘pseudo-bond’ connecting *p*′ and *q*′ A normal ‘pinch’ **f** has a force dipole directed along the **r**_*p*′_ − **r**_*q*′_ line (the **n**_*p*′*q*′_ direction). Such a pinch is expected to induce a hinge motion. A shear ‘pinch’ will be in a perpendicular direction ⊥ **n**_*p*′*q*′_, and is expected to induce a shear motion.

In the protein scenario, we try to tune the spring network to exhibit a low-energy mode in which the protein is divided into two sub-domains moving like rigid bodies. This large-scale mode can be detected by examining the relative motion of two neighboring vertices, *p* and *q* at another location at the boundary (usually at the opposite side). Such a desired response at the other side of the protein is specified by a response vector **v**, whose only non-zero entries correspond to the directions of the response at *p* and *q*. Again, we usually consider a ‘dipole’ response **v**_*q*_ = −**v**_*p*_.

### Evolution and Mutation

The quality of the response, *i.e*., the biological fitness, is specified by how well the response follows the desired one **v**. In the context of our model, we chose the (scalar) observable *F* as *F* = **v**^t^**u** = **v**_*p*_**u**_*p*_ + **v**_*q*_**u**_*q*_ = **v**^t^**Gf** [2]. In an evolution simulation one would exchange AAs between H and P while demanding that the fitness change *F* is positive or non-negative. By this, we mean *F >* 0 is thanks to a beneficial mutation, whereas *δF* = 0 corresponds to a neutral one. Deleterious mutations *δF <* 0 are generally rejected. We may impose a stricter minimum condition, *δF ≥ *ε* F* with a small positive *ε*, say 1%. An alternative, stricter criterion would be the demand that each of the terms in *F*, **v**_*p*_**u**_*p*_ and **v**_*q*_**u**_*q*_, increases separately.

### Evolving the Green function using Dyson’s and Woodbury’s formulas

Dyson’s formula follows from the identity 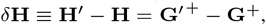, which is multiplied by **G** on the left and **G**′ on the right to yield [6]. The formula remains valid for the pseudo-inverses in the non-singular sub-space. One can calculate the change in fitness by evaluating the effect of a mutation on the Green function, **G**′ = **G** + *δ***G**, and then examining the change, *δF* = **v**^t^*δ***Gf** [3]. Using [6] to calculate the mutated Green function **G**′ is an impractical method as it amounts to inverting at each step a large *n_d_×n_d_* matrix. However, the mutation of an AA at *i* has a localized effect. It may change only up to *z* = 12 bonds among the bonds *α*(*i*) with the neighboring AAs. Thanks to the localized nature of the mutation, the corresponding defect Hamiltonian *δ***H**_*i*_ is therefore of a small rank, *r* ≤ *z* = 12, equal to the number of switched bonds (the average *r* is about 9.3). *δ***H**_*i*_ can be decomposed into a product *δ***H**_*i*_ = **MBM**^t^. The diagonal *r×r* matrix **B** records whether a bond *α*(*i*) is switched from weak to strong (**B**_*αα*_ = *k_s_ − k_w_* = +0.99) or vice versa (**B**_*αα*_ = −0.99), and **M** is a *n_d_×r* matrix whose *r* columns are the bond vectors **m**_*α*_ for the switched bonds *α*(*i*). This allows one to calculate changes in the Green function more efficiently using the Woodbury formula (61, 62), 
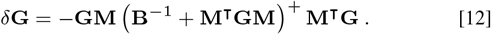

The two expressions for the mutation impact *δ***G**, [6] and [12], are equivalent and one may get the scattering series of [6] by a series expansion of the pseudo-inverse in [12]. The practical advantage of [12] is that the only (pseudo-) inversion it requires is of a low-rank tensor (the term in brackets). This accelerates our simulations by a factor of (*n_a_/r*)^3^ ≃ 10^4^.

### Pathologies and broken networks

A network broken into disjoint components exhibits floppy modes owing to the low energies of the relative motion of the components with respect to each other. The evolutionary search might end up in such non-functional unintended modes. The common pathologies we observed are: (i) isolated nodes at the boundary that become weakly connected via H→P mutations, (ii) ‘sideways’ channels that terminate outside the target region (which typically includes around 8−10 sites), and (iii) channels that start and end at the target region without connecting to the binding site. All these are some easy-to-understand floppy modes, which can vibrate independently of the location of the pinch and cause the response to diverge (>*F*_m_) without producing a functional mode. We avoid such pathologies by applying the pinch force to the protein network symmetrically: pinch the binding site on face L and look at responses on face R and vice versa. Thereby we not only look for the transmission of the pinch from the left to right but also from right to left. The basic algorithm is modified to accept a mutation only if it does not weaken the two-way responses and enables hinge motion of the protein. This prevents the vibrations from being localized at isolated sites or unwanted channels.

### Dimension and Singular Value Decomposition (SVD)

To examine the geometry of the fitness landscape and the genotype-to-phenotype map, we looked at the correlation among numerous solutions, typically *N*_sol_ ∼ 10^6^. Each solution is characterized by three vectors: (i) the gene of the functional protein, **c**_*_, (a vector of length *n_a_* = 200 codons) (ii) the flow field (displacement), **u**(**c**_*_) = **G**(**c**_*_)**f**, (a vector of length *n_d_* = 400 of *x* and *y* velocity components), (iii) the shear field **s**_*_ (a vector of length *n_a_* = 200). We compute the shear as the symmetrized derivative of the displacement field using the method of (18, 32). The values of the **s**_*_ field is the sum of squares of the traceless part of the strain tensor (Frobenius norm). These three types of vectors are stored along the rows of three matrices *W_C_*, *W_U_* and *W_S_*. We calculate the eigenvectors of these matrices, **C**_*k*_, **U**_*k*_ and **S**_*k*_, via singular value decomposition (SVD) (as in (23)). The corresponding SVD eigenvalues are the square roots of the eigenvalues of the covariance matrix *W*^t^*W*, while the eigenvectors are the same. In typical spectra, most eigenvalues reside in a continuum bulk region that resembles the spectra of random matrices. A few larger outliers, typically around a dozen or so, carry the non-random correlation information.

### The protein backbone

A question may arise as to what extent the protein’s backbone might affect the results described so far. Proteins are polypeptides, linear heteropolymers of AAs, linked by covalent peptide bonds, which form the protein backbone. The peptide bonds are much stronger than the non-covalent interactions among the AAs and do not change when the protein mutates. We therefore augmented our model with a ‘backbone’: a linear path of conserved strong bonds that passes once through all AAs. We focused on two extreme cases, a serpentine backbone either parallel to the shear band or perpendicular to it (Fig. 6).

**Fig. 6.**
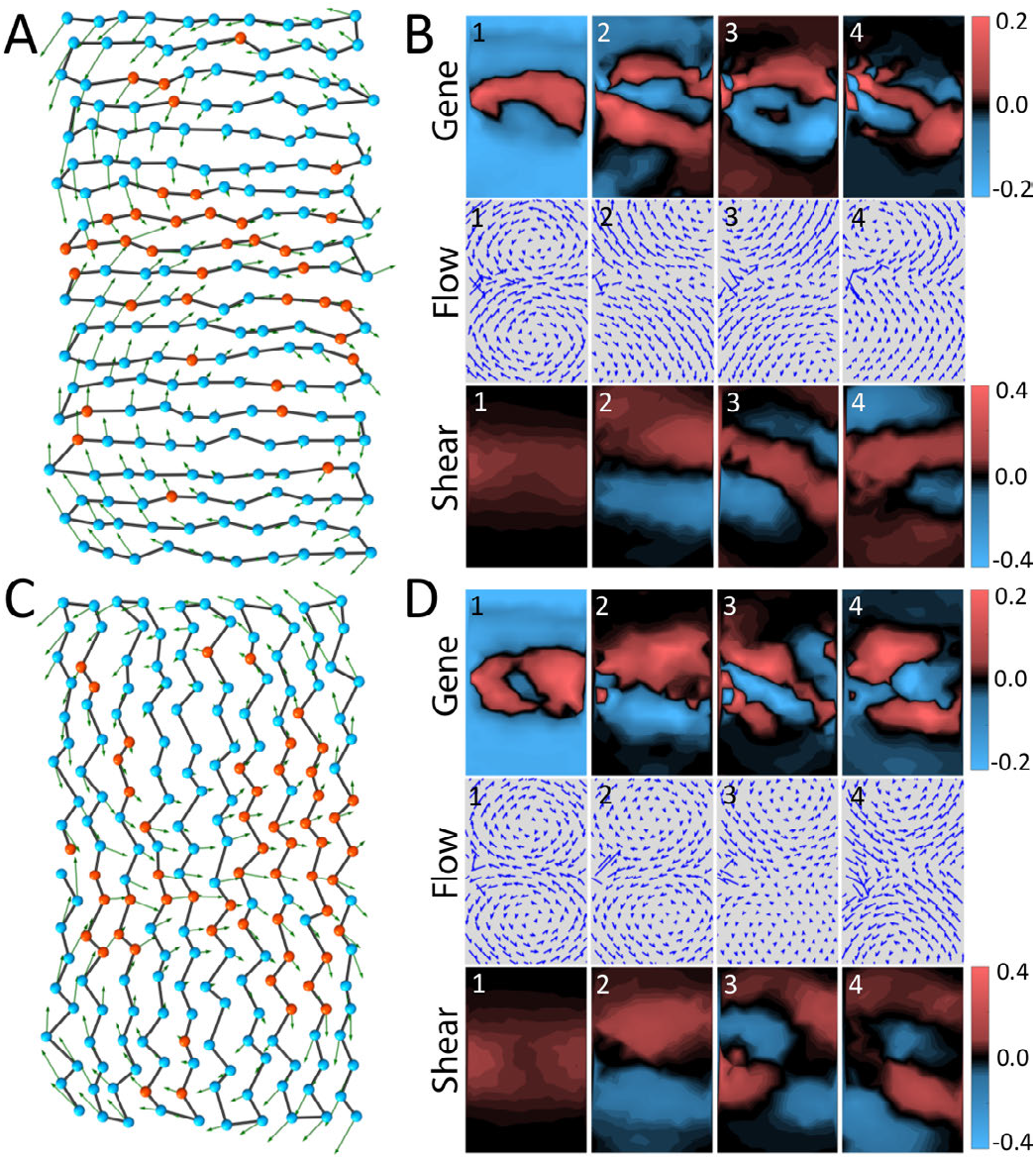
The effect of the backbone on evolution of mechanical function. The backbone induces long-range mechanical correlations which influence protein evolution. We examine two configurations: parallel (A-B) and perpendicular to the channel (C-D). Parallel: (A) The backbone directs the formation of a narrow channel along the fold (compared to Fig. 5A). (B) First four SVD eigenvectors of the gene **C**_*k*_ (top row), the flow **U**_*k*_ (middle) and the shear **S**_*k*_ (bottom). Perpendicular: (C) The formation of the channel is ‘dispersed’ by the backbone. (D). First four SVD eigenvectors of **C**_*k*_ (top row), **U**_*k*_ (middle) and shear **S**_*k*_ (bottom).

The presence of the backbone does not interfere with the emergence of a low-energy mode of the protein whose flow pattern (*i.e*., displacement field) is similar to the backbone-less case with two eddies moving in a hinge like fashion. In the parallel configuration, the backbone constrains the channel formation to progress along the fold (Fig. 6A). The resulting channel is narrower than in the model without backbone (Figs. 1D, 5). In the perpendicular configuration, the evolutionary progression of the channel is much less oriented (Fig. 6C). While the flow patterns are similar, closer inspection shows noticeable differences, as can be seen in the flow eigenvectors **U**_*k*_ (Fig. 6B,D). The shear eigenvectors **S**_*k*_ represent the derivative of the flow, and therefore highlight more distinctly these differences.

As for the correspondence between gene eigenvectors **C**_*k*_ and shear eigenvectors **S**_*k*_, the backbone affects the shape of the channel in concert with the sequence correlations around it. Transmission of mechanical signals appears to be easier along the orientation of the fold (parallel configuration, Fig. 6A). Transmission across the fold (perpendicular configuration) necessitates significant deformation of the backbone and leads to ‘dispersion’ of the signal at the output (Fig. 6C). We propose that the shear band will be roughly oriented with the direction of the fold, but this requires further analysis of structural data. Overall, we conclude from our examination that the backbone adds certain features to patterns of the field and sequence correlation, without changing the basic results of our model. The presence of the backbone might constrain the evolutionary search, but this has no significant effect on the fast convergence of the search and on the long-range correlations among solutions. We expect that in realistic 3D geometry the backbone will have a weaker effect than what we observed in 2D networks, since the extra dimension adds more options to avoid the backbone constraint.

## ACKNOWLEDGMENTS

We thank Jacques Rougemont for calculations of shear in glucokinase (Fig. 1B) and for helpful discussions. We thank Stanislas Leibler, Michael R. Mitchell, Elisha Moses, Giovanni Zocchi and Olivier Rivoire for helpful discussions and encouragement. We thank Steve Granick and Alex Petroff for valuable comments on the manuscript. JPE is supported by an ERC advanced grant ‘Bridges’, and TT by the Institute for Basic Science IBS-R020 and the Simons Center for Systems Biology of the Institute for Advanced Study, Princeton.

## References

1. Daniel RM, Dunn RV, Finney JL, Smith JC (2003) The role of dynamics in enzyme activity. Annu Rev Biophys Biomol Struct 32:69–92.

2. Bustamante C, Chemla YR, Forde NR, Izhaky D (2004) Mechanical processes in biochemistry. Annual Review of Biochemistry 73:705–748.

3. Hammes-Schiffer S, Benkovic SJ (2006) Relating protein motion to catalysis. Annual Review of Biochemistry 75:519–541.

4. Boehr DD, McElheny D, Dyson HJ, Wright PE (2006) The dynamic energy landscape of dihydrofolate reductase catalysis. Science 313(5793):1638–1642.

5. Koonin EV, Wolf YI, Karev GP (2002) The structure of the protein universe and genome evolution. Nature 420(6912):218–223.

6. Zeldovich KB, Shakhnovich EI (2008) Understanding protein evolution: from protein physics to darwinian selection. Annu Rev Phys Chem 59:105–127.

7. Povolotskaya IS, Kondrashov FA (2010) Sequence space and the ongoing expansion of the protein universe. Nature 465(7300):922–926.

8. Liberles DA, et al. (2012) The interface of protein structure, protein biophysics, and molecular evolution. Protein Sci 21(6):769–785.

9. Dill KA, MacCallum JL (2012) The protein-folding problem, 50 years on. Science 338(6110):1042–1046.

10. Perutz MF (1970) Stereochemistry of cooperative effects in haemoglobin: Haem-haem interaction and the problem of allostery. Nature 228(5273):726–734.

11. Henzler-Wildman KA, et al. (2007) Intrinsic motions along an enzymatic reaction trajectory. Nature 450(7171):838–U13.

12. Huse M, Kuriyan J (2002) The conformational plasticity of protein kinases. Cell 109(3):275–282.

13. Savir Y, Tlusty T (2010) Reca-mediated homology search as a nearly optimal signal detection system. Mol Cell 40(3):388–396.

14. Goodey NM, Benkovic SJ (2008) Allosteric regulation and catalysis emerge via a common route. Nat Chem Biol 4(8):474–482.

15. Koshland D (1958) Application of a theory of enzyme specificity to protein synthesis. Proc Natl Acad Sci U S A 44(2):98–104.

16. Savir Y, Tlusty T (2007) Conformational proofreading: the impact of conformational changes on the specificity of molecular recognition. PLoS One 2:e468.

17. Gerstein M, Lesk AM, Chothia C (1994) Structural mechanisms for domain movements in proteins. Biochemistry 33(22):6739–6749.

18. Mitchell MR, Tlusty T, Leibler S (2016) Strain analysis of protein structures and low dimensionality of mechanical allosteric couplings. Proc Natl Acad Sci U S A 113(40):E5847–E5855.

19. Mitchell MR, Leibler S (2017) Elastic strain and twist analysis of protein structural data and allostery of the transmembrane channel kcsa. Physical Biology.

20. Qu H, Zocchi G (2013) How enzymes work: A look through the perspective of molecular viscoelastic properties. Phys Rev X 3(1).

21. Joseph C, Tseng CY, Zocchi G, Tlusty T (2014) Asymmetric effect of mechanical stress on the forward and reverse reaction catalyzed by an enzyme. PLoS One 9(7).

22. Tlusty T (2016) Self-referring dna and protein: a remark on physical and geometrical aspects. Phil. Trans. Roy. Soc. A 374(2063).

23. Tlusty T, Libchaber A, Eckmann JP (2017) Physical model of the genotype-to-phenotype map of proteins. Phys Rev X 7(2).

24. Cui Q, Karplus M (2008) Allostery and cooperativity revisited. Protein Science 17(8):1295–1307.

25. Daily MD, Upadhyaya TJ, Gray JJ (2008) Contact rearrangements form coupled networks from local motions in allosteric proteins. Proteins-Structure Function and Bioinformatics 71(1):455–466.

26. Motlagh HN, Wrabl JO, Li J, Hilser VJ (2014) The ensemble nature of allostery. Nature 508(7496):331–339.

27. Hemery M, Rivoire O (2015) Evolution of sparsity and modularity in a model of protein allostery. Physical Review E 91(4).

28. Flechsig H (2017) Design of elastic networks with evolutionary optimized long-range communication as mechanical models of allosteric proteins. Biophysical Journal 113(3):558–571.

29. Rocks JW, et al. (2017) Designing allostery-inspired response in mechanical networks. Proc Nat Acad Sci USA 114(10):2520–2525.

30. Yan L, Ravasio R, Brito C, Wyart M (2017) Architecture and coevolution of allosteric materials. Proc Nat Acad Sci USA 114(10):2526–2531.

31. Miyashita O, Onuchic JN, Wolynes PG (2003) Nonlinear elasticity, proteinquakes, and the energy landscapes of functional transitions in proteins. Proc Natl Acad Sci U S A 100(22):12570–12575.

32. Gullett PM, Horstemeyer MF, Baskes MI, Fang H (2008) A deformation gradient tensor and strain tensors for atomistic simulations. Modell Simul Mater Sci Eng 16(1).

33. Dill KA (1985) Theory for the folding and stability of globular proteins. Biochemistry 24(6):1501–1509.

34. Green G (1828) An essay on the application of mathematical analysis to the theories of electricity and magnetism. (Printed for the author, by T. Wheelhouse, Nottingham).

35. Abrikosov A, Gorkov L, Dzyaloshinski I (1963) Methods of Quantum Field Theory in Statistical Physics. (Parentice Hall).

36. Cordell HJ (2002) Epistasis: what it means, what it doesn’t mean, and statistical methods to detect it in humans. Human Molecular Genetics 11(20):2463–2468.

37. Phillips PC (2008) Epistasis - the essential role of gene interactions in the structure and evolution of genetic systems. Nat Rev Genet 9(11):855–867.

38. Mackay TFC (2014) Epistasis and quantitative traits: using model organisms to study gene-gene interactions. Nat Rev Genet 15(1):22–33.

39. Ortlund EA, Bridgham JT, Redinbo MR, Thornton JW (2007) Crystal structure of an ancient protein: Evolution by conformational epistasis. Science 317(5844):1544–1548.

40. Breen MS, Kemena C, Vlasov PK, Notredame C, Kondrashov FA (2012) Epistasis as the primary factor in molecular evolution. Nature 490(7421):535–538.

41. Gong LI, Suchard MA, Bloom JD (2013) Stability-mediated epistasis constrains the evolution of an influenza protein. Elife 2:e00631.

42. Miton CM, Tokuriki N (2016) How mutational epistasis impairs predictability in protein evolution and design. Protein Sci 25(7):1260–1272.

43. Tirion MM (1996) Large amplitude elastic motions in proteins from a single-parameter, atomic analysis. Phys Rev Lett 77(9):1905–1908.

44. Bahar I, Rader AJ (2005) Coarse-grained normal mode analysis in structural biology. Curr Opin Struct Biol 15(5):586–592.

45. Chennubhotla C, Rader AJ, Yang LW, Bahar I (2005) Elastic network models for understanding biomolecular machinery: From enzymes to supramolecular assemblies. Phys Biol 2(4):S173–S180.

46. Penrose R (1955) A generalized inverse for matrices. Mathematical Proceedings of the Cambridge Philosophical Society 51(3):406–413.

47. Ben-Israel A, Greville TN (2003) Generalized inverses: theory and applications. (Springer Science & Business Media) Vol. 15.

48. Tewary VK (1973) Green-function method for lattice statics. Advances in Physics 22(6):757–810.

49. Elliott RJ, Krumhansl JA, Leath PL (1974) The theory and properties of randomly disordered crystals and related physical systems. Rev. Mod. Phys. 46(3):465–543.

50. Dyson FJ (1949) The *s* matrix in quantum electrodynamics. Phys. Rev. 75(11):1736–1755.

51. Desai MM, Weissman D, Feldman MW (2007) Evolution can favor antagonistic epistasis. Genetics 177(2):1001.

52. Göbel U, Sander C, Schneider R, Valencia A (1994) Correlated mutations and residue contacts in proteins. Proteins: Structure, Function, and Bioinformatics 18(4):309–317.

53. Te¸sileanu T, Colwell LJ, Leibler S (2015) Protein sectors: Statistical coupling analysis versus conservation. PLoS Comput Biol 11(2):e1004091.

54. Savir Y, Noor E, Milo R, Tlusty T (2010) Cross-species analysis traces adaptation of rubisco toward optimality in a low-dimensional landscape. Proc Natl Acad Sci U S A 107(8):3475–3480.

55. Savir Y, Tlusty T (2013) The ribosome as an optimal decoder: A lesson in molecular recognition. Cell 153(2):471–479.

56. Kaneko K, Furusawa C, Yomo T (2015) Universal relationship in gene-expression changes for cells in steady-growth state. Phys. Rev. X 5(1):011014.

57. Friedlander T, Mayo AE, Tlusty T, Alon U (2015) Evolution of bow-tie architectures in biology. PLOS Computational Biology 11(3):e1004055.

58. Miyazawa S, Jernigan RL (1985) Estimation of effective interresidue contact energies from protein crystal structures: quasi-chemical approximation. Macromolecules 18(3):534–552.

59. Born M, Huang K (1954) Dynamical theory of crystal lattices, The International series of monographs on physics. (Clarendon Press, Oxford,), pp. xii, 420 p.

60. Alexander S (1998) Amorphous solids: Their structure, lattice dynamics and elasticity. Phys Rep 296(2–4):65–236.

61. Woodbury MA (1950) Inverting modified matrices, Statistical Research Group, Memo. Rep. no. 42. (Princeton University, Princeton, N. J.), p. 4.

62. Deng CY (2011) A generalization of the sherman–morrison–woodbury formula. Applied Mathematics Letters 24(9):1561–1564.

